# Biochemical characterization and NMR study of a PET-hydrolyzing cutinase from *Fusarium solani pisi*

**DOI:** 10.1101/2022.11.01.514593

**Authors:** Kristina Naasen Hellesnes, Shunmathi Vijayaraj, Peter Fojan, Evamaria Petersen, Gaston Courtade

**Author notes:** These authors contributed equally (Kristina Naasen Hellesnes, Shunmathi Vijayaraj).

## Abstract

In recent years, the drawbacks of plastics have become evident, with plastic pollution becoming a major environmental issue. There is an urgent need to find solutions to efficiently manage plastic waste by using novel recycling methods. Biocatalytic recycling of plastics by using enzyme-catalyzed hydrolysis is one such solution that has gained interest, in particular for recycling polyethylene terephthalate (PET). To provide insights into PET hydrolysis by cutinases, we have here characterized the kinetics of a PET-hydrolyzing cutinase from *Fusarium solani pisi* (FsC) at different pH values, mapped the interaction between FsC and the PET analog BHET by using NMR spectroscopy, and monitored product release directly and in real time by using time-resolved NMR experiments. We found that primarily aliphatic side chains around the active site participate in the interaction with BHET, and that pH conditions and mutation around the active site (L182A) can be used to tune the relative amounts of degradation products. Moreover, we propose that the low catalytic performance of FsC on PET is caused by poor substrate binding combined with slow MHET hydrolysis. Overall, our results provide insights into obstacles that preclude efficient PET hydrolysis by FsC and suggest future approaches for overcoming these obstacles and generating efficient PET-hydrolyzing enzymes.

**TOC Graphic (For Table of Contents use only):** 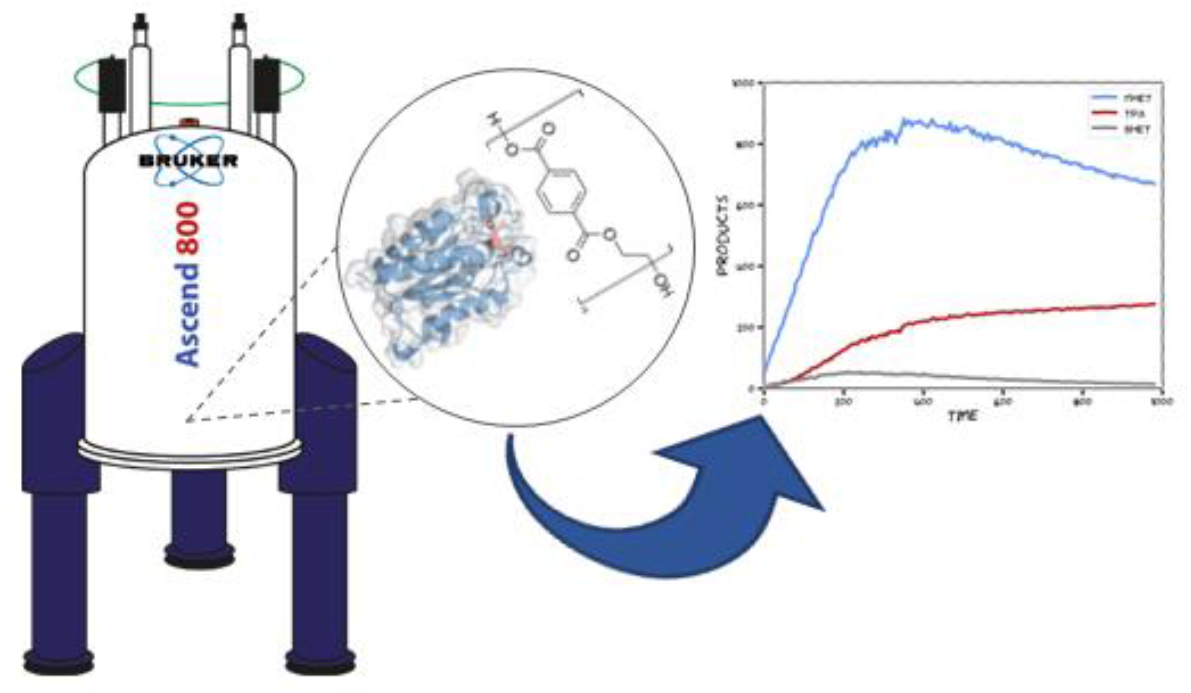

## INTRODUCTION

Enzymatic depolymerization of polyethylene terephthalate (PET) by cutinases (EC 3.1.1.74)^1,2^ and cutinase-like PETases (EC 3.1.1.101)^3,4^ have recently received enormous attention because of their ability to hydrolyze the scissile ester bonds is PET, yielding well-defined products (BHET: bis(2-hydroxyethyl) terephthalate; MHET: mono(2-hydroxyethyl) terephthalic acid; TPA: terephthalic acid; EG: ethylene glycol) that can be reused to make new plastics.^1,5^ These enzymes have thus provided a novel alternative to thermomechanical recycling of plastics, a process in which only clear, homogenous plastic can be recycled with quality loss in each cycle (i.e., downcycling)^6^.

Important enzymes for PET hydrolysis include a PETase from *Ideonella sakaiensis* (IsP)^2–4^, and cutinases from *Thermobifida fusca* (TfC)^2,7,8^, *Humicola insolens* (HiC)^2,9,10^, *Fusarium solani pisi* (FsC)^9,11^, and leaf-branch compost cutinase (LCC)^1^. Cutinases are serine esterases that possess a Ser-His-Asp catalytic triad^12^. They have a characteristic α/β-hydrolase fold and naturally hydrolyze ester bonds in cutin, an insoluble polyester in plant cuticle composed of hydroxy and epoxy fatty acids^13^.

With increasing implementation of enzymes in plastic recycling processes, a “polyester biorefinery” may be envisioned in which hydrolysates from PET feed stocks can be used for different recycling and upcycling applications^14^. In this context, it would be desirable to not only increase the catalytic efficiency^15^ and thermostability^16^ of PET hydrolases, but also understand how reaction conditions influence product distribution. Moreover, overcoming factors limiting the catalytic efficiency of the enzymes is a requirement for their efficient use.

In order to shine light on these issues, we have used a combination of NMR spectroscopy and UV-based assays to characterize FsC (UniProtKB: P00590). Using continuous time-resolved NMR experiments we followed the hydrolysis of PET by FsC under different pD values, and used the backbone amide resonances to probe the interaction of an inactive S120A-FsC mutant with BHET. Moreover, we applied a suspension-based assay^17^ to derive inverse Michaelis-Menten kinetic parameters for FsC. Overall, our results provide useful biochemical insights into PET hydrolysis by FsC.

## MATERIALS AND METHODS

### Particle size measurement

The particle size distribution of the PET powder was measured with a Mastersizer 3000 Hydro MV (Malvern) instrument. Approximately 0.5 g of PET powder was dissolved in 10 mL 96% ethanol. Solutions were added to the cell dropwise until an obscuration of 4% was obtained. Refractive indices of 1.636 and 1.360 were used for PET and ethanol, respectively, and a particle absorption index of 0.010 was used. The data from five measurements was analyzed using the Mastersizer software, which provides average particle size parameters (volume mean diameter and surface mean diameter), as well as the specific surface area of the particles.

### Protein production and purification

Recombinant *E. coli* ER2566 cells (New England Biolabs T7 Express) harboring the pFCEX1D plasmid (containing wild-type FsC, S120A-FsC or L182A-FsC) were incubated in 5 mL precultures containing LB supplemented with 100 µg/mL ampicillin at 30 °C and 225 rpm for 16 hours. Main cultures were made by inoculating 500 mL of either 2xLB (20 g/L tryptone, 10 g/L yeast extract, 5 g/L NaCl) or ^15^N-enriched minimal M9 medium (6 g/L Na_2_HPO_4_, 3 g/L KH_2_PO_4_, 0.5 g/L NaCl supplemented with 98% (^15^NH_4_)_2_SO_4_, 4 g/L D-(+)-glucose, 5 mL Gibco MEM Vitamin Solution (100x), 300 mg/mL MgSO_4_, 2 mg/L ZnSO_4_, 10 mg/L FeSO_4_, 2 mg/L CuSO_4_, and 20 mg/L CaCl_2_) with 1% preculture, followed by incubation at 25 °C and 225 rpm. At OD_600_ = 1.7 – 1.9, the cells were induced with 0.1 mM isopropyl-β-D-thiogalactopyranoside followed by further incubation at 25 °C and 225 rpm overnight.

Cells were harvested by centrifugation for 5 min at 5000 g and 4 °C, and periplasmic fractions were prepared by the osmotic shock method as follows. The pellet was gently resuspended on ice in 50 mL TES buffer (100 mM Tris HCl, 500 mM sucrose, 0.5 mM ethylenediaminetetraacetic acid (EDTA), pH 7.5) with a cOmplete ULTRA protease inhibitor tablet (Roche). After 10 min centrifugation at 6500 g, the pellet was resuspended on ice in 50 mL ultrapure water. The suspension was then centrifuged for 15 min at 15000 g followed by 30 min at 21000 g. The TES and water fractions were dialyzed at 4 °C in 2 L reverse-osmosis water overnight. Equilibration buffer (25 mM Na-acetate pH 5.0) was added to both fractions followed by centrifugation at 7000 g and 4 °C for 5 min. The supernatant was filtered using a filter (0.2 µm pore size) prior to further protein purification.

The proteins were purified by loading the periplasmic extracts in a 20 mM Na-acetate buffer pH 5.0 onto a 5 mL HiTrap CM FF cation exchanger (Cytiva) connected to an ÄKTApure FPLC system (Cytiva). All steps were performed at a flow rate of 5 mL/min. Proteins were eluted by using a linear salt gradient (0 – 500 mM NaCl). FsC and S120A-FsC eluted at 40 – 120 mM NaCl. Eluted fractions were analyzed using sodium dodecyl sulphate-polyacrylamide gel electrophoresis (SDS-PAGE; Figure S1) gels run under denaturing conditions using SurePAGE Bis-Tris gels (GenScript) and MES-SDS running buffer (GenScript) followed by staining using the eStain L1 Protein Staining System (GenScript). PAGE-MASTER Protein Standard Plus (GenScript) was used for the identification of target proteins.

The fractions containing wild-type FsC, S120A-FsC or L182A-FsC were pooled and concentrated using centrifugal concentrators (10 kDa cut-off, Sartorius). The protein concentration was calculated by measuring A_280_ using Nanodrop and the theoretical extinction coefficient (ε = 14690 M^-1^ cm^-1^), which was estimated using the ProtParam server (https://web.expasy.org/protparam/)^18^. The yields were calculated to be approximately 40 mg protein per L cell culture.

### Interactions with BHET

Interactions between S120A-FsC and BHET were probed by measuring chemical shift perturbations (CSP) as follows. A ^15^N-HSQC spectrum of ^15^N-labeled S120A-FsC (175 µM) in a buffer consisting of 25 mM phosphate pH 5 and 10 mM NaCl with 10% D2O, was recorded at 313K as a reference. BHET was dissolved in another sample of ^15^N-S120A-FsC (175 µM), and the two samples were combined in different proportions to obtain the following BHET concentrations: 0.3 mM, 1.1 mM, 3.6 mM, 5.5 mM, and 7.6 mM while keeping the protein concentration constant. ^15^N-HSQC spectra were recorded for each BHET concentration. CSP in amide pairs were expressed as the combined chemical shift change, 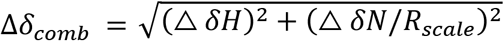. where Δ*δH* and Δ*δN* are the CSP of the amide proton and nitrogen, respectively, and *R*_*scale*_ was set 6.5^19^. The dissociation constant, *K*_*D*_, was calculated by fitting CSP to a two-site fast exchange model, 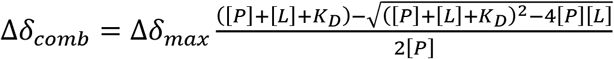. where Δ*δ*_*max*_ is the CSP at full saturation, and [*P*] and [*L*] are respectively the concentration of S120A-FsC and BHET.

These NMR spectra were recorded in a Bruker Ascend 600 MHz spectrometer equipped with an Avance III HD console and a 5-mm cryogenic CP-TCI z-gradient probe at the NV-NMR laboratory at NTNU.

### Suspension-based assay

The kinetics of the FsC reaction on PET were measured by using a suspension-based assay originally described by Bååth et al^17^, at three different pH values (5.0, 6.5 and 9.0). Reactions were set up in triplicate in Eppendorf tubes with a total volume of 600 µL, containing 10 g L^-1^ crystalline PET powder (GoodFellow product code ES306031; 37.7 ± 2.6% crystallinity^20^), enzyme concentrations varying between 0 – 1 µM, and either a 25 mM sodium acetate buffer pH 5.0, a 25 mM sodium phosphate buffer pH 6.5 containing 50 mM NaCl, or a 50 mM glycine buffer pH 9.0.

The reactions were incubated in an Eppendorf ThermoMixer C at 40 °C and 450 rpm for 7 hours. At 0, 1, 3, 5 and 7 hours 100 µL of were transferred from each reaction to a 96-well MultiScreen_HTS_ HV Filter Plate (0.45 µm pore size; Millipore), and the reactions were stopped by vacuum filtering using a Vac-Man 96 Vaccum Manifold (Promega) onto a 96-well Clear Flat Bottom UV-Transparent Microplate (Corning). PET hydrolysis products were quantified by measuring A_240_ in a Spectramax Plus 284 microplate reader (Molecular Devices) and concentrations were calculated by using a standard curve made with 15, 30, 60, 90, 120 and 150 µM TPA (Figure S2).

### Time-resolved ^1^H-NMR experiments

Time-resolved ^1^H-NMR experiments were carried out in a Bruker Avance III HD 800 MHz spectrometer equipped with a 5-mm cryogenic CP-TCI z-gradient probe at the NV-NMR laboratory at NTNU.

The buffers used were the same as for the suspension-based assay, but they were lyophilized and redissolved in 99.9% D_2_O (pD 5.0) prior to use, giving pD values at 5.0, 6.5, and 9.0. Reactions (600 µL) were prepared in 5 mm NMR tubes and contained an amorphous PET film (GoodFellow product code ES301445; 2.0 ± 1.6% crystallinity^20^) cut into a size of 30×4×0.25 mm, buffer, FsC (10 µM) and TSP (trimethylsilylpropanoic acid; 400 µM).

After adding wild-type FsC or FsC-L182A, samples were immediately inserted into the spectrometer, where they were incubated for 17.5 hours at 40 °C. A solvent-suppressed ^1^H spectrum was acquired every 5 min by using a modified version of the 1D NOESY pulse sequence with presaturation and spoil gradients (noesygppr1d). Briefly, a 2D matrix was made with the direct dimension (TD2 = 32k) corresponding to the 1D ^1^H experiment spectrum, and the indirect dimension (TD1 = 196) corresponding to number of individual experiments. The experiment time was determined by the acquisition time (AQ = 1.7 s), number of scans (NS = 32), the NOESY mixing time (D8 = 10 ms), the relaxation delay (D1 = 4 s), and an interexperiment delay (D14 = 130 s).

Signals corresponding to the aromatic protons of BHET, MHET and TPA were integrated using the serial integration (intser) routine in Bruker TopSpin version 4.1.3.

## RESULTS AND DISCUSSION

### Particle size distribution of PET powder

Enzymatic activity on PET is affected by the physical properties of the substrate, like percent crystallinity, particle size, and accessible surface area^20^. To provide more information about commonly used PET substrates we used laser diffraction to measure the particle size distribution (volume-weighted mean diameter, D[4,3] = 103 ± 1 µm; surface area-weighted mean diameter, D[3,2] = 65.3 ± 0.7 µm) and specific surface area (92 ± 1 mm^2^ mg^-1^) of crystalline PET powder (Figure S3).

### Interactions between S120A-FsC and BHET

To identify the substrate-binding residues on FsC we titrated BHET, as an analog of PET, on the inactive S120A-FsC mutant and followed chemical shift perturbations (CSP) on the amide proton-nitrogen pairs by using ^15^N-HSQC spectra. Upon substrate binding, changes in the chemical environment around protein nuclei cause corresponding changes in ^15^N-HSQC signals. The previously published^21^ chemical shift assignment of FsC (Biological Magnetic Resonance Data Bank (BMRB) accession 4101) was used for analysis of ^15^N-HSQC data.

Addition of BHET to ^15^N-labeled S120A-FsC led to gradual changes in the ^1^H-^15^N resonances consistent with fast exchange between the free and bound states^22^. Analysis of CSP allows estimation of dissociation constants, but interpretation of CSP with a small Δd_*max*_ (A120 in Figure 1A) can lead unreliable estimates. Analyzing CSP with higher Δd_*max*_ values on residues near the active site (Figure 1A) led to an estimate of around *K*_*D*_ = 10 – 20 mM. This is a very weak interaction and, as discussed below, it may be one of the reasons for the low catalytic activity of FsC. However, suitable estimation of *K*_*D*_ values requires full saturation of the protein, which was unreachable due to the poor solubility of BHET^22^.

**Figure 1.**
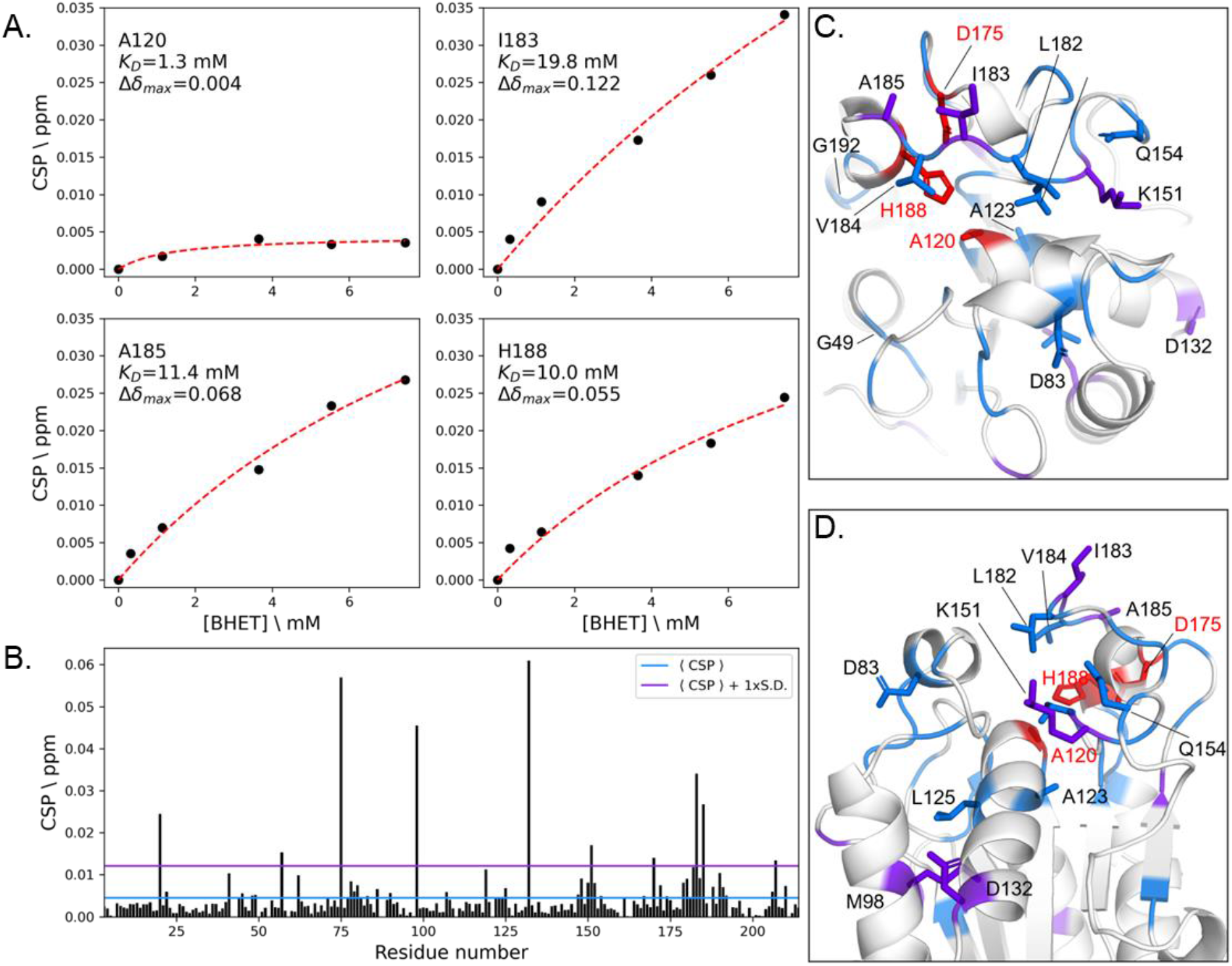
Interactions between S120A-FsC and BHET. Panel A show chemical shift perturbations (CSP; black dots) at increasing BHET concentrations for four representative residues near the active site. The dissociation constant (*K*_*D*_) and maximum CSP (Δ*δ*_*max*_) are derived from the fit of the data to a two-site fast exchange model (red line). Panel B shows the CSP per residue; residues with CSP larger than the average CSP, ⟨*CSP* ⟩, are colored blue in Panels C and D, whereas residues with CSP larger than the average CSP plus one standard deviation are colored purple in Panels F and G. Residues in the active site are colored red in Panels C and D.

Titration with 7.6 mM BHET led to CSP (Figure 1B) mainly on residues located around the active site of FsC (S120A, D175 and H188), where several aliphatic residues (A123, L125, L182, I183, V184, A185), as well as some polar residues (D83, T150, K151, Q154) were affected by the interaction with BHET (Figure 1C–D). This suggests that the binding is predominantly mediated by hydrophobic interactions. CSP on more distant residues (M98, D132) are likely the result of structural rearrangements upon binding with BHET, rather than direct interactions.

There are similarities between these findings and those of a recent study in which Charlier et al^23^ used NMR was to probe binding of LCC to MHET. Regions around LCC’s V212–A213 (equivalent to L182–V184 in FsC) and H191 (equivalent to K151 in FsC) were also found to be important for binding MHET, but based on our results (Figure 1C–D) BHET binding seems to require a more extended binding pocket in the regions around G49 and G192.

### Effect of pH on FsC-catalyzed PET hydrolysis

The electrostatic potential inside and around the active site of cutinases has been hypothesized to be closely linked to catalytic efficiency^24^. To test this hypothesis, we assayed the enzymatic activity of FsC on PET powder at different pH values and enzyme concentrations, and analyzed the data by fitting an inverse Michaelis-Menten model^2^ that has previously been used to characterize cutinase hydrolysis of PET. In contrast to the conventional Michaelis-Menten model in which *V*_*max*_*/E*_*0*_ describes the catalytic rate when all enzymes are substrate-bound, the inverse model uses ^inv^*V*_*max*_/*S*_*0*_ to define the rate when all sites on the insoluble PET substrate are saturated with enzymes (for a detailed description of these models in the context of PET degradation see Bååth et al.^2,25^). The model described the data well (Figure 2, Table 1), and maximum activity in terms of ^inv^*V*_*max*_/*S*_*0*_ was found at pH 9.0. At this alkali pH, the concentration of solubilized products was approximately 3-fold higher than at pH 5, and 1.5-fold higher than at pH 6.5. This observation is consistent with previous reports on the “electrostatic catapult” mechanism of esterases and lipases^26^, where electrostatic repulsion (favored by high pH) of negatively charged hydrolysis products (MHET and TPA in the case of PET) from negative charges in the active site cleft favors catalytic performance. Reduction in pH was accompanied by a decrease in enzymatic activity together with an increase in binding affinity (i.e., a reduction of ^inv^*K*_*M*_ values) (Table 1). This observation finds explanation in the neutralization of negative charges on the substrate, hydrolysis products and active site, which reduce electrostatic repulsion effects^24^. This may lead to too tight binding of the enzyme to substrate and/or products, precluding efficient catalysis. The validity of this interpretation hinges on the assumption that ^inv^*K*_*M*_ can be used as a proxy to describe enzyme-substrate affinity.

**Table 1.** Inverse Michaelis-Menten parameters for FsC on PET powder at 40 °C and three pH values. The parameters were calculated based on fitting of the data in Figure 2. Catalytic rates are given with two units, ^inv^*V*_*max*_/*S*_*0*_ in nmol products per gram PET powder per second, and ^inv^*V*_*max*_/*S*_*0*_*\** in nmol products per particle surface area in m^2^ per second. The error bars represent standard error from the fit (n=3).

| pH | $^{inv}K_M$ (μM) | $^{inv}V_{max}/S_0$ (nmol g<sup>-1</sup> s<sup>-1</sup>) | $^{inv}V_{max}/S_0^*$ (nmol m<sup>-2</sup> s<sup>-1</sup>) |
| --- | --- | --- | --- |
| 5.0 | 0.15 ± 0.09 | 0.23 ± 0.03 | 2.5 ± 0.3 |
| 6.5 | 0.46 ± 0.13 | 0.49 ± 0.07 | 5.3 ± 0.8 |
| 9.0 | 0.57 ± 0.18 | 0.74 ± 0.09 | 8.0 ± 0.9 |

**Figure 2.**
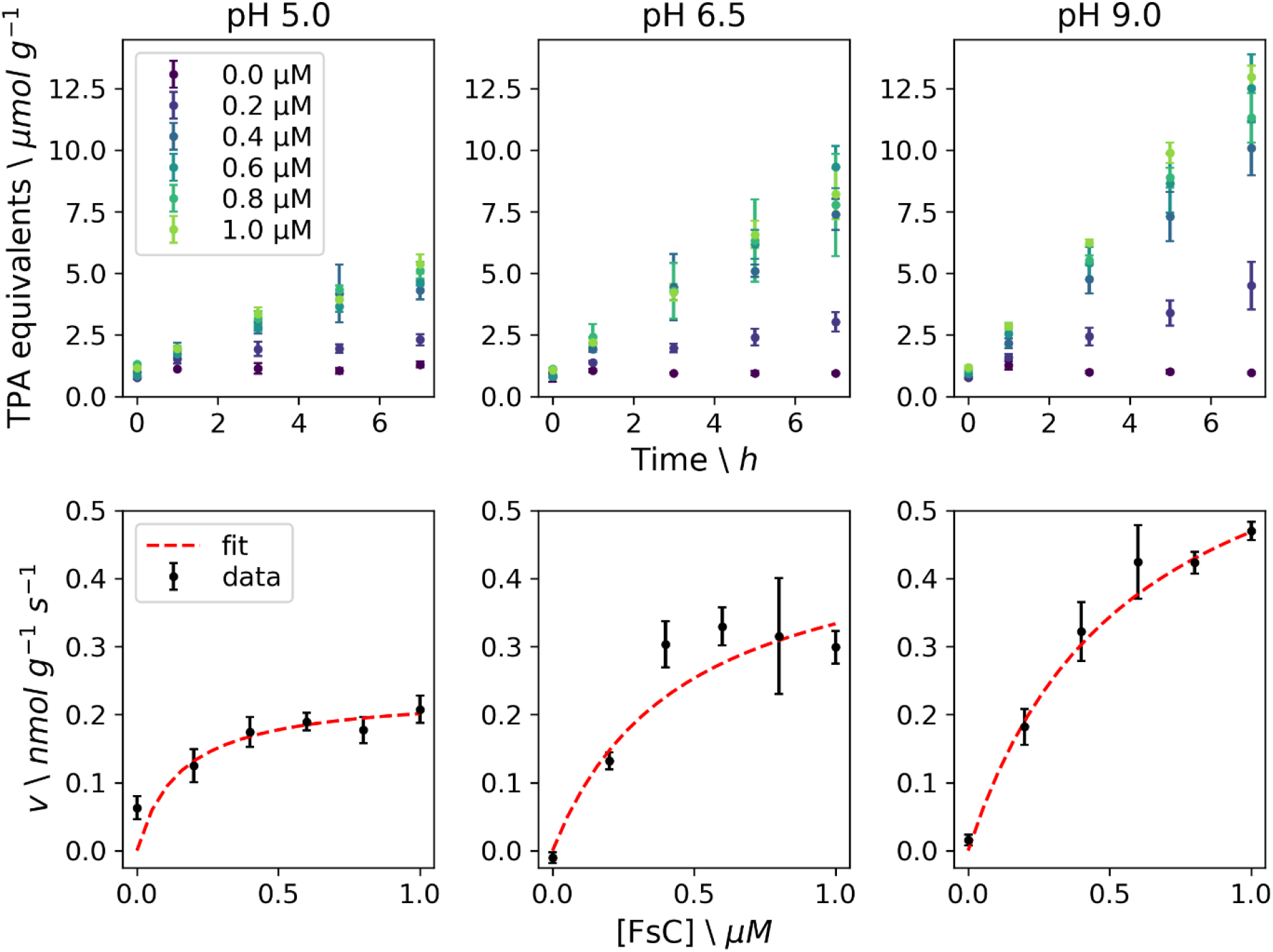
Enzymatic activity for wild-type FsC on PET powder (10 g L^-1^) at 40 °C and three pH values. The top panels show the release of TPA equivalents per gram PET powder with increasing enzyme concentration (0 – 1 µM) at 0, 1, 3, 5 and 7 hours. The bottom panels show the initial rate, *v* (based on a linear fit of the product concentration at 0, 1 and 3 hours), as a function of enzyme concentration (black dots), and the corresponding fit of the inverse Michaelis-Menten model (red dashed line). The error bars correspond to the standard deviation (n = 3).

Interestingly, the ^inv^*K*_*M*_ values here reported (Table 1) match the ^inv^*K*_*M*_ values found by Bååth et al^25^ for TfC and LCC in the presence of surfactant, resulting in maximum ^inv^*V*_*max*_/*S*_*0*_ values of about 9 (TfC) and 40 (LCC) nmol g^-1^ s^-1^. However, these values are 10 – 100-fold higher than the ^inv^*V*_*max*_/*S*_*0*_ values for FsC (Table 1). The inferior performance of FsC on PET may be caused by its poor binding to BHET (Figure 1) and PET (similar to the surfactant-weakened binding affinities of TfC and LCC). A structure-based sequence alignment (Figure S4) of FsC (PDB 1CEX) to TfC (PDB 5ZOA) and LCC (PDB 4EB0) reveals that FsC has a 3_10_-helix (L81–R88; η2 in Figure S4) in its active site cleft, which is absent in TfC and LCC. This helix participates at least via D83 in the interaction of the enzyme with BHET (Figure 1). It may be that the presence of this helix is detrimental for the binding and catalytic activity of FsC on PET. Araújo et al^27^ have previously shown that L81A (also part of the η2 helix) and L182A FsC mutants had higher hydrolytic activity on PET and polyamide 6,6 fibers than the wild-type cutinase, indicating that engineering a less crowded binding site could boost cutinase activity on PET.

### Hydrolytic activity on PET films monitored by time-resolved NMR

Suspension-based assays on microplates require manual sampling over long time periods to obtain kinetic data. This drawback of discontinuous assays has recently been addressed by the development of a continuous UV-based assay^28^. Here we demonstrate the applicability of a continuous assay based on time-resolved NMR spectroscopy. An advantage of time-resolved NMR is that the technique allows direct observation of all intermediates and products simultaneously (Figure S5), providing direct insights into the reaction progress. However, NMR signals can be affected by other factors than product formation, such as line broadening due to inhomogeneities in the magnetic field caused by the presence of insoluble substrate in the NMR tube. Even though caution should be taken when comparing NMR-derived activity profiles between samples, trends can be appreciated in the activity profiles (Figure 3). In all conditions only lesser amounts of BHET were seen in the activity profiles, suggesting that BHET is hydrolyzed at a faster rate than PET. The main product from PET hydrolysis at all pD values is MHET, which comprises about 80% of the products (Figure 3; WT bottom panels). After about 400 minutes, at pD 6.5 and 9.0, the concentration of MHET decreases linearly at a slower rate and is accompanied by an increase in the TPA concentration. The slower hydrolysis of MHET hydrolysis to TPA suggests that MHET is not a preferred substrate for FsC.

**Figure 3.**
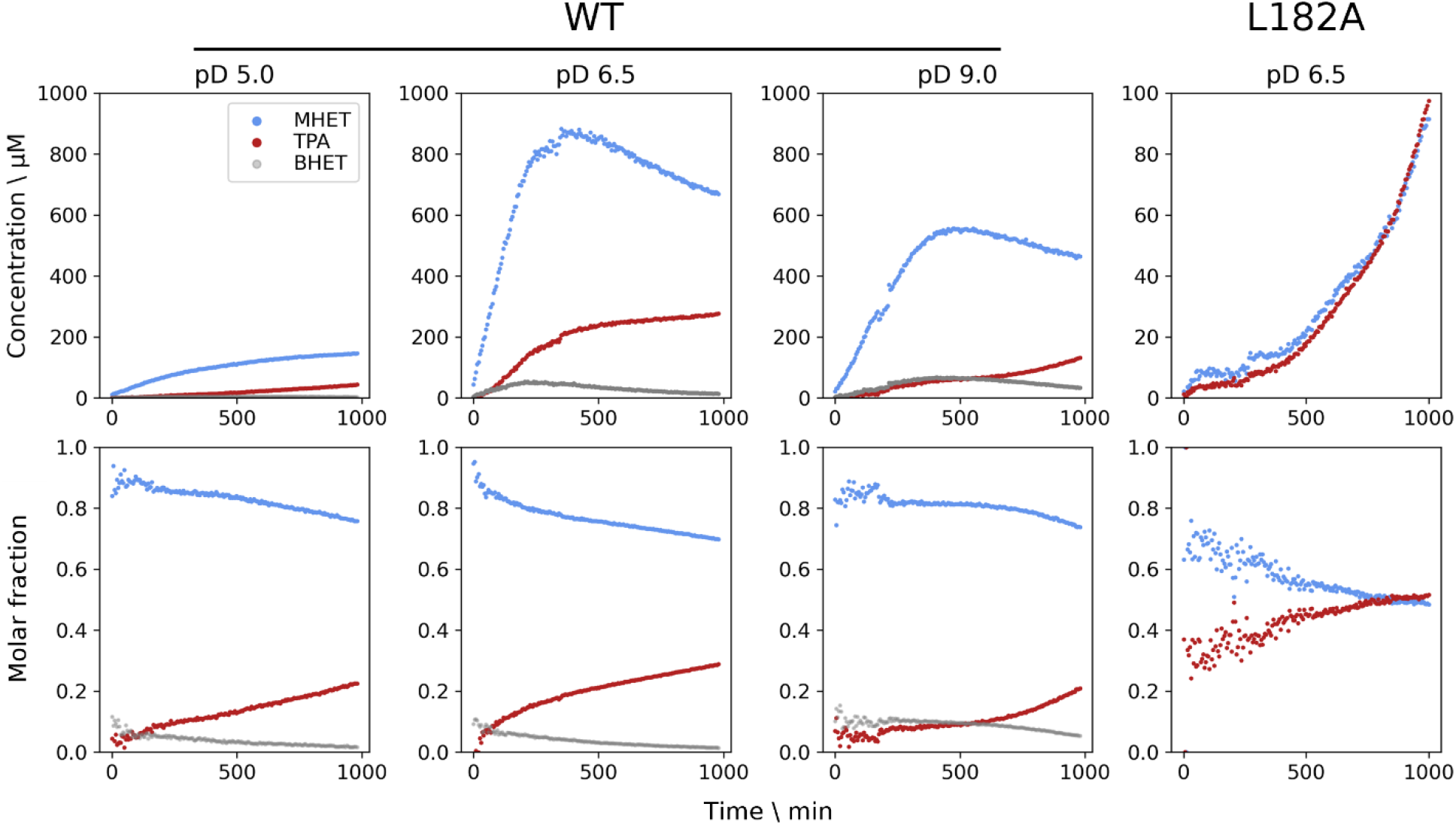
Enzymatic activity measured by time-resolved ^1^H NMR for wild-type FsC and L182A-FsC on a PET film at 40 °C and different pD values. The profiles for each product are proportional to the integrals of the aromatic proton signals, monitored via 196 individual ^1^H spectra recorded every 5 min for a total of 16.3 hours. The top panels show the product concentration over time, which was calculated based on the integral ratio to a TSP signal (corresponding to 400 µM) used as an internal standard. The bottom panels show the molar fraction of the products. The structures and chemical shift assignments of the degradation products are shown in Figure S5. Note that the y-scale for the top panel of L182A is different.

Using the same time-resolved NMR method we investigated a FsC-L182A mutant that has previously^27^ been shown to have higher PET hydrolyzing activity. The time courses (Figure 3; L182A panels) show production of approximately equal amounts of both TPA and MHET, a markedly different product distribution to the wild-type enzyme. However, in contrast to earlier findings^27^ (see above), the product yields after 1000 minutes were about 8-fold lower than for the wild-type enzyme. We propose the following reasoning for the difference in relative yields. Whereas we detected TPA and MHET hydrolysis products directly and by using an enzyme concentration of 10 µM, the experimental conditions used by Araújo et al were markedly different. The authors used approximately a 100-fold higher enzyme concentration, and a detection method in which only TPA is observed indirectly via fluorescence detection of hydroxyterephthalate, which is produced by reacting TPA with hydroxy radicals at 90°C^29^. This means that only TPA amounts were detected and, since FsC-L182A produces more TPA than the wild-type enzyme (Figure 3), it is possible that the overall catalytic performance of the mutant was overestimated by Araujo et al. Moreover, the different enzyme loadings, which are known to significantly affect the catalytic performance of PET hydrolyzing enzymes^20^, could have further contributed to the discrepant yields.

At pD 9.0, the decrease in MHET (and increase in TPA) concentration after 400 minutes appears to be slower than at the other pD values (Figure 3). This suggests that, in addition to mutations around the active site, the pH and pD conditions may be used to tune the relative amounts of degradation products, which could be of interest for optimizing enzymatic synthesis of MHET by cutinases^30^.

In contrast with the PET powder assay where pH 9.0 gave highest activity, time-resolved NMR assays on PET films at different pD values showed that pD 6.5 (and not pD 9.0) resulted in maximum enzymatic activity (Figure 3). Ronkvist et al^9^ have previously observed that FsC activity on PET varies little from pH 6.5 to 8.5, but it drops sharply at pH 9.0. This observation has previously been observed to agree with the pH range where FsC is most stable; its maximum thermostability is found at pH 6 – 8.5 but it decreases sharply at pH values outside the range^24^. Differences in optimal pH and pD values for maximum enzymatic activity measurements on PET powder and PET films may thus be explained by FsC having lower thermal stability in the assays with PET films. The PET powder has both higher crystallinity (37.7 ± 2.6%^20^) and surface area (552 mm^2^) than PET films (crystallinity: 2.0 ± 1.6%^20^; surface area: 257 mm^2^). These morphological differences likely translate to variations in protein-substrate interactions, which influence enzymatic activity.

## CONCLUSION

We have characterized a PET-hydrolyzing cutinase from *F. solani pisi*, FsC, by using a combination of NMR spectroscopy and kinetic studies at different pH and pD values. In summary, our results show that continuous time-resolved NMR experiments are a useful tool to assay enzymatic activity on PET, complementing discontinuous UV-based plate assays. These assays we show that pD conditions and an amino acid around the active site (i.e., L182) influence product distribution (i.e. the TPA-to-MHET ratio), and that weak interactions between FsC and BHET/PET, combined with inefficient hydrolysis of MHET, likely contribute to the lower catalytic activity of FsC on PET compared to other cutinases (e.g., TfC and LCC). NMR titration experiments providing insights into the molecular interaction of FsC with BHET can be used for future studies seeking to engineer FsC for use in biocatalytic plastic recycling applications.

## DATA AVAILABILITY

Data and python scripts used for data processing and making figures are available from https://github.com/gcourtade/papers/tree/master/2023/FsC-PET.

## ACCESSION CODES

*Fusarium solani pisi* cutinase (FsC): P00590

## ACKNOWLEDGMENT

G.C. was funded by the Novo Nordisk Foundation via the project number

NNF18OC0032242.We would like to thank Dr. Morten J. Dille for technical assistance.

