## Supplementary Information for "Biochemical characterization and NMR study of a PET-hydrolyzing cutinase from *Fusarium solani pisi*"

[‡]*These authors contributed equally.*

**SUPPLEMENTARY INFORMATION**

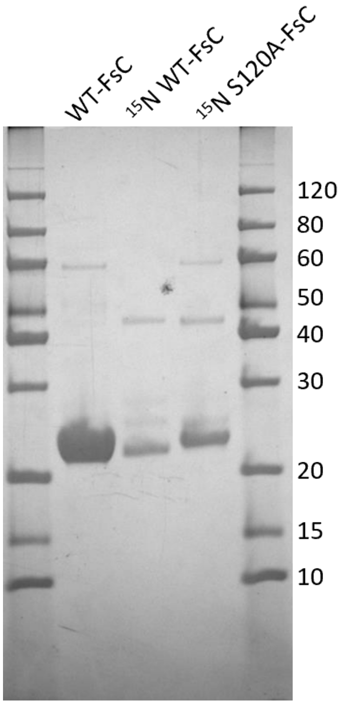


**Figure S1. Sodium dodecyl sulphate-polyacrylamide gel electrophoresis (SDS-PAGE).** The picture of the stained gel shows the FsC samples used for activity assays (WT-FsC) and NMR titration experiments (15N S120A-FsC), as well as a sample of 15N WT-FsC. The molecular weights (in kDa) of the standards (PAGE-MASTER Protein Standard Plus from GenScript) are indicated for the bands in right-most standard lane.

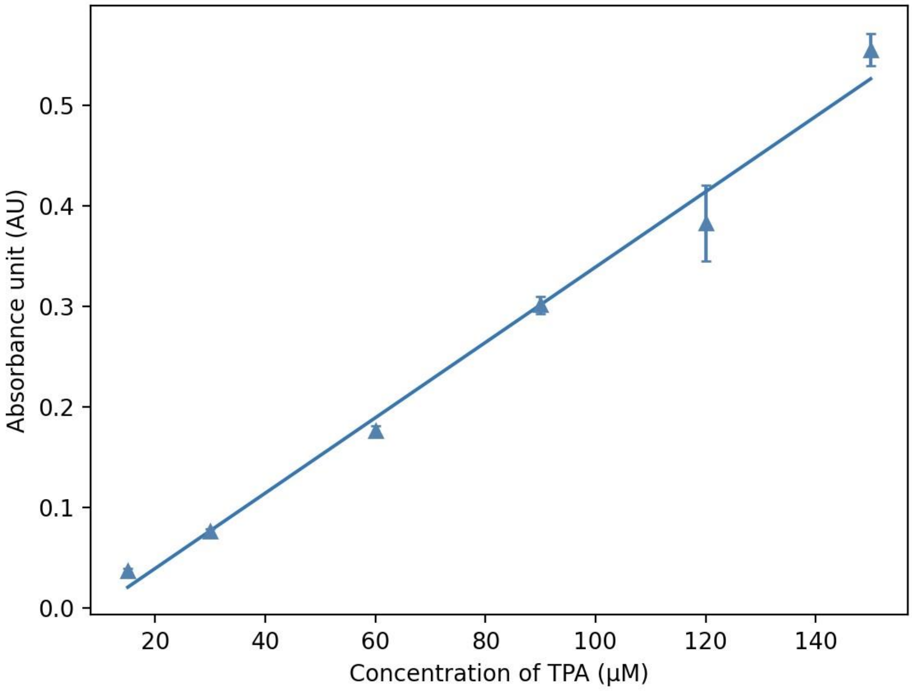


**Figure S2. Terephthalic acid (TPA) standard curve.** Measured absorbance values at 240 nm as a function of TPA concentration. Linear regression was performed and gave a linear function of $A_{240}$ = 0.0038*[TPA] - 0.0359, with $R^2$ = 0.99.

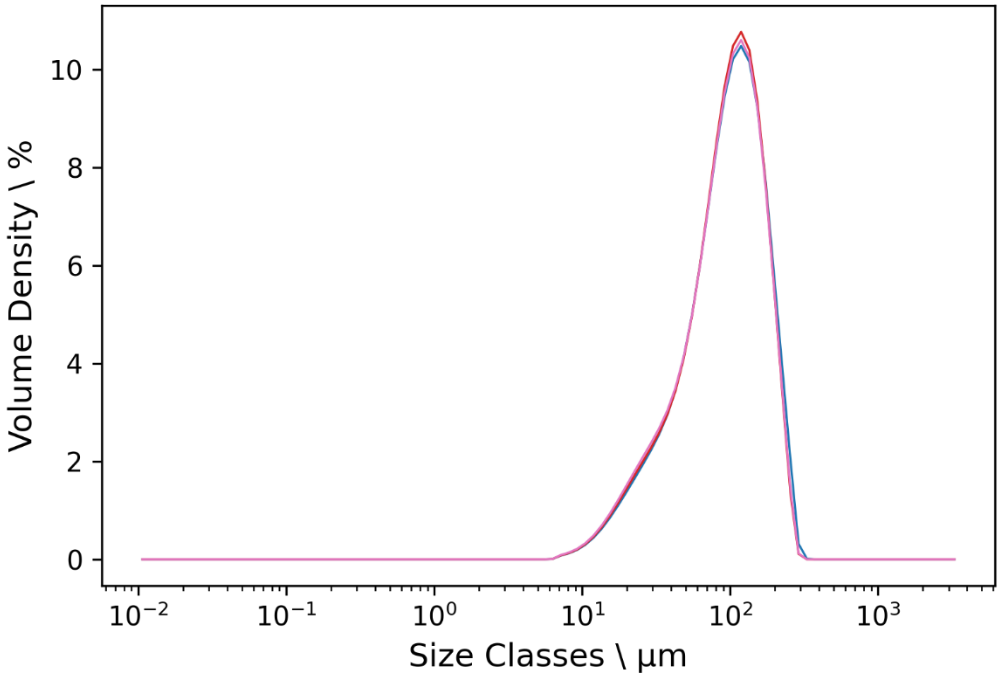


**Figure S3. Analysis of particle size distribution of PET powder.** The figure shows on overall of five curves describing the distribution of weight classes in a sample of crystalline PET powder (GoodFellow product code ES306031) dispersed in 96% ethanol. The following particle size parameters were determined. Volume-weighted mean diameter, D[4,3] = 103 ± 1 μm; surface area-weighted mean diameter, D[3,2] = 65.3 ± 0.7 μm and specific surface area = 92 ± 1 $mm^2$ $mg^{-1}$.

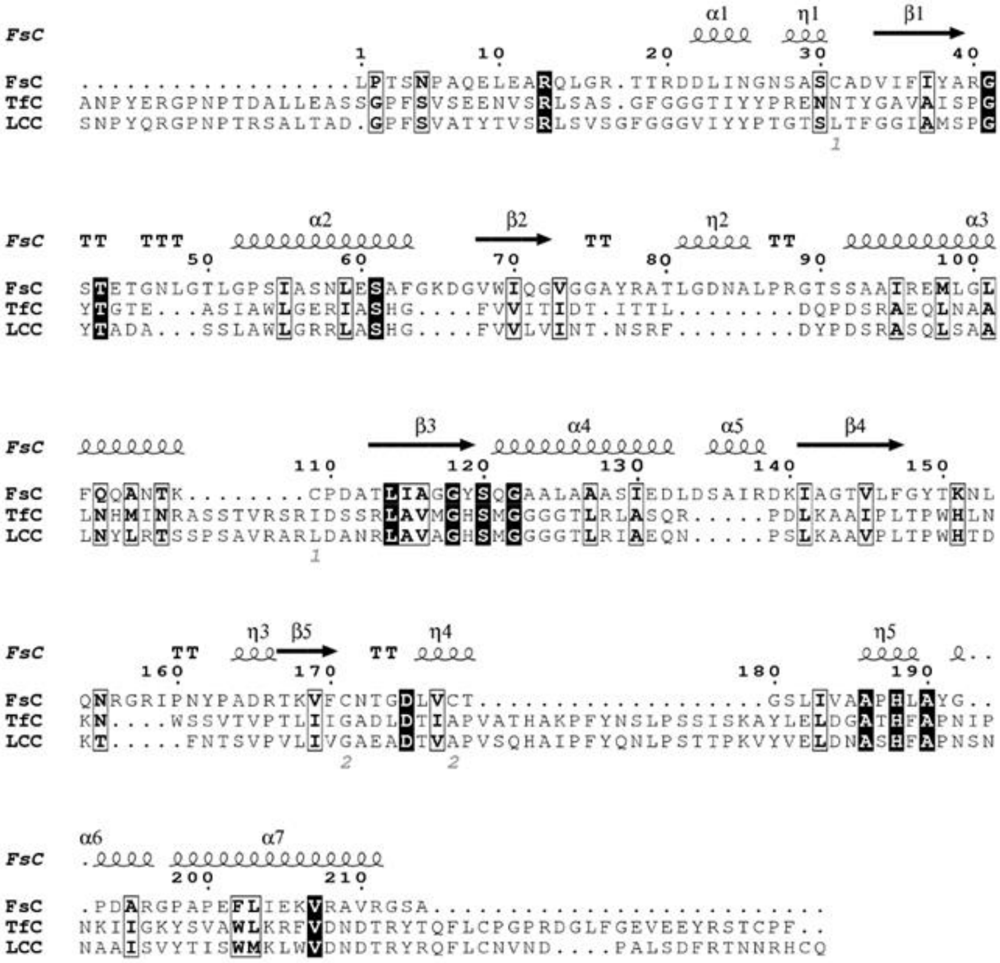

**Figure S4. Multiple-sequence alignment (MSA) of three cutinases.** The MSA was performed in the Expresso/T-coffee server[1] and the visual representation was made using the ESPript 3.0 server[2]. FsC (*Fusarium solani pisi* cutinase; PDB 1CEX), TfC (*Thermobifida fusca* cutinase; PDB 5ZOA), LCC (leaf-branch compost cutinase; PDB 4EB0). Secondary structure elements from FsC are shown on top (helices with squiggles, β-strands with arrows and turns with TT letters). The bottom numbers (1 and 2) represent the disulphide bonds in FsC. α and η represent alpha- and $3_{10}$-helices, respectively. Identical and similar residues are highlighted black and boxed white, respectively.

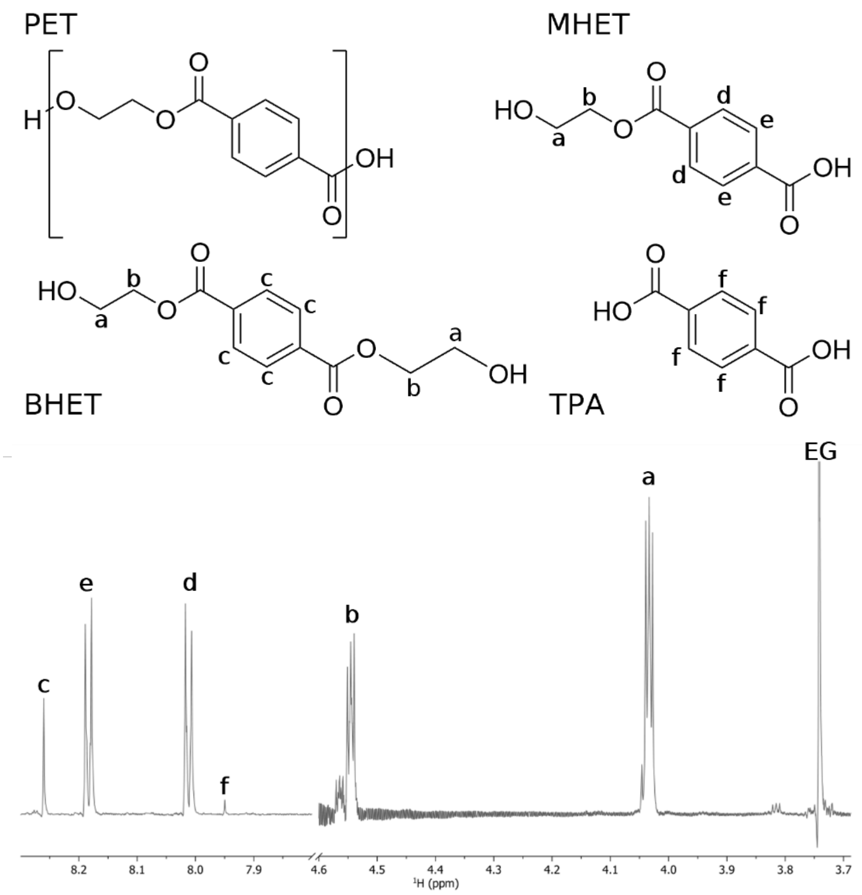


**Figure S5. Chemical structures and chemical shifts.** The structures of polyethylene terephthalate (PET), bis(2-hydroxyethyl) terephthalate (BHET), mono(2-hydroxyethyl) terephthalic acid (MHET), and terephthalic acid (TPA) are shown. The chemical shifts corresponding to $^{1}H$ and $^{13}C$ resonances (in ppm) were assigned at pD 6.5 and 313 K as **a**: 4.02, 59.8; **b**: 4.53, 63.8; **c:** 8.19,129.5; **d:** 8.01, 128.8; **e:** 8.18, 129.5; **f:** 7.94, 129.5. The 1H NMR spectrum under the structures shows an overview of the assigned resonances, as well as the free ethylene glycol (EG) peak at 3.74 ppm.
